# Integrative Machine Learning Reveals Potential Signature Genes Using Transcriptomics in Colon Cancer

**DOI:** 10.1101/2025.02.28.640917

**Authors:** Mostafa Amir Hamza, Md. Saiful Islam

## Abstract

**Background:** Colon cancer is a significant health burden in the world and the second leading cause of cancer-related deaths. Despite advancements in diagnosis and treatment, identifying robust biomarkers for early detection and therapeutic targets remains imperative.

**Materials and methods:** This study used an integrative approach combining transcriptomics and machine learning to identify signature genes and pathways associated with colon cancer. RNA-Seq data from The Cancer Genome Atlas-Colon Adenocarcinoma (TCGA-COAD) project, comprising 485 samples (444 tumors and 41 normal tissues), were analyzed.

**Results:** Differential gene expression analysis revealed 657 upregulated and 8,566 downregulated genes. Notably, EPB41L3, TSPAN7, and ABI3BP were identified as highly upregulated, while LYVE1, PLPP1, and NFE2L3 were significantly downregulated in tumor samples. Gene Set Enrichment Analysis (GSEA) identified dysregulated pathways, including E2F targets, MYC targets, and G2M checkpoints, underscoring cell cycle regulation and metabolic reprogramming alterations in colon cancer. Machine learning models-Random Forest, Neural Networks, and Logistic Regression-achieved high classification accuracy (97–99%) and near-perfect Area Under the Receiver Operating Characteristic Curve (AUC-ROC) values (approximately 1.00), validating their predictive capabilities. Key genes consistently identified across these models highlight their potential translational relevance as biomarkers. This study integrates differential expression analysis, pathway enrichment, and machine learning to uncover critical insights into colon cancer biology.

**Conclusion:** The findings lay the groundwork for developing diagnostic and therapeutic strategies, with the identified genes and pathways serving as promising candidates for future validation and clinical applications. This approach exemplifies the potential of precision medicine to advance colon cancer research and improve patient outcomes.

## Introduction

Colon cancer is a major global health concern, ranking as the third most common cancer worldwide and accounting for approximately 10% of all cancer cases. It is also the second leading cause of cancer-related deaths globally. Despite advancements in diagnosis and treatment, the lack of robust biomarkers for early detection and therapeutic targeting remains a critical challenge. In 2020, an estimated 1.9 million new cases of colorectal cancer and over 930,000 related deaths were reported worldwide^1-3^. This highlights the urgent need for innovative approaches to uncover molecular drivers of colon cancer and translate them into clinical applications.

Excluding skin cancers, colon cancer is one of the most frequently diagnosed cancers in both men and women. It ranks third in cancer-related deaths among men and fourth among women, but collectively, it is the second leading cause of cancer mortality. In 2024, the American Cancer Society projects approximately 106,590 new cases of colon cancer (54,210 in men and 52,380 in women)^4^. Alarmingly, the incidence of colon cancer is increasing among younger adults, where it has become the leading cause of cancer-related deaths in men under 50 and the second leading cause in women under 50, following breast cancer. Each generation born since the 1950s faces a higher risk than the previous one. Colon cancer is expected to account for a significant proportion of the 53,010 CRC-related deaths anticipated in the United States in 2024^5, 6^.

These statistics underscore the urgent need for regular screening, early detection, and lifestyle modifications to mitigate the risk of colon cancer. Advancements in RNA sequencing (RNA-Seq) have revolutionized transcriptomics, enabling comprehensive profiling of gene expression across various biological conditions, including cancer^7^. The Cancer Genome Atlas (TCGA) provides publicly available RNA-Seq datasets that facilitate in-depth molecular analyses of multiple cancer types^8^.

This study focuses on identifying differentially expressed genes (DEGs) and pathways in colon cancer, a malignancy characterized by significant clinical challenges and heterogeneity. Differential expression analysis serves as a foundational step in uncovering genes with altered expression in tumors, while Gene Set Enrichment Analysis (GSEA) offers a pathway-level understanding of systemic changes in tumor biology^9^.

Additionally, gene interaction networks provide a systems biology perspective, revealing key nodes and hubs that may serve as regulatory elements or therapeutic targets^10^. With the growing emphasis on precision medicine, the integration of machine learning with transcriptomics has gained momentum. Machine learning approaches, such as logistic regression, artificial neural networks, and random forests, offer powerful tools for feature selection, pattern recognition, and predictive modeling^11^. In this study, we leveraged these techniques using Python’s sci-kit-learn library to identify potential marker genes capable of distinguishing tumors from normal samples, intending to enhance diagnostic and prognostic capabilities in colon cancer. By integrating these diverse methodologies, this study provides a holistic approach to biomarker discovery in colon cancer. It identifies potential marker genes and establishes a framework for leveraging RNA-Seq data in translational cancer research. The findings hold promise for advancing our understanding of colon cancer biology and contributing to developing personalized diagnostic and therapeutic strategies.

## Materials and Methods

### Data Acquisition and Preprocessing

Colon adenocarcinoma (COAD) gene expression data were retrieved from the TCGA database (https://portal.gdc.cancer.gov/repository) using TCGAbiolinks in R. We analyzed 485 samples (444 tumors and 41 normal tissues) after filtering for clinical data availability. Transcriptomic data from the TCGA-COAD project were selected, comprising 485 out of 524 samples: 444 tumor tissue samples and 41 matched normal tissue samples based on clinical data availability. Clinical data for 459 colon cancer patients were also downloaded, with key survival and staging information extracted for analysis.

### Gene Expression Pre-Filtering and Normalization

Using TCGAbiolinks, transcriptomics data were downloaded as fragments per kilobase million (FPKM) unstranded normalized data^12, 13^. The dataset initially included 60,660 Ensembl gene identifiers. These Ensembl IDs were converted to gene names, resulting in 42,225 named genes. Genes with zero expression values across all samples were excluded, leaving 10,962 genes for further expression and machine learning analysis. Since the data was normalized, a log2 transformation was applied to the FPKM data for consistency.

### Gene Expression Analysis

Log2-transformed normalized data were used to assess gene expression alterations in tumor patients compared to those in healthy controls. A t-test was performed to identify significant gene expression changes, followed by false discovery rate (FDR) correction to adjust for multiple comparisons. Differentially expressed genes were defined as those with an FDR-adjusted p-value < 0.05. These genes were visualized using a heatmap and volcano plot. Sample clustering patterns were examined through a Principal Component Analysis (PCA) plot to explore group separations.

### Gene Set Enrichment Analysis (GSEA)

Based on the differentially expressed genes (p-value < 0.05), hallmark pathway analysis was performed using the Molecular Signatures Database (MSigDB) with GSEA. The identified pathways were visualized using GSEA plots to understand the gene sets associated with specific pathway enrichment patterns^14, 15^.

### Building the Machine Learning Model

Machine learning models, including Random Forest (RF), Neural Network (NN), and Logistic Regression (LR), were implemented using the Scikit-learn (sklearn) package in Python to analyze the filtered TCGA RNA-seq dataset^16-18^. The dataset was preprocessed by excluding genes with zero expression values across all samples, resulting in 10,962 genes, and a log2 transformation was applied to the normalized FPKM data. To ensure robust training and evaluation, the dataset was split into training (70%) and testing (30%) sets using stratified sampling to preserve the proportion of tumor and normal samples.

The Random Forest model was constructed using an ensemble learning approach with 100 decision trees, utilizing Gini impurity as the splitting criterion. Hyperparameters such as maximum tree depth and the minimum number of samples required for splits were optimized through grid search and cross-validation. The Neural Network model was designed as a multilayer perceptron with three hidden layers, each consisting of 50 neurons, and employed the rectified linear unit (ReLU) activation function. The Adam optimizer was used for weight updates, and early stopping was applied to mitigate overfitting. Hyperparameters, including the learning rate and batch size, were fine-tuned to maximize performance. The Logistic Regression model employed L2 regularization to prevent overfitting, with the maximum number of iterations set to 1,000 to ensure model convergence.

Model performance was assessed using accuracy, area under the receiver operating characteristic curve (AUC), and F1-score as evaluation metrics. The predictions generated by each model were further analyzed to identify significant marker genes associated with tumor and normal samples, providing insights into their predictive relevance and biological significance.

## Results

### Differential Gene Expression Analysis

After pre-filtering and normalization, we identified 10,962 genes with non-zero expression values suitable for further analysis (**Supplementary Table 1**). Differential expression analysis revealed 657 upregulated and 8,566 downregulated genes in tumor samples compared to matched normal controls (p-value < 0.05). Notably, EPB41L3, TSPAN7, and ABI3BP were among the most upregulated genes, while LYVE1, PLPP1, and NFE2L3 were significantly downregulated. A heatmap of the differentially expressed genes clearly demonstrated distinct segregation between tumor and normal samples, underscoring significant transcriptomic differences (**Fig. 1A, Supplementary Table 2 and 3**). Principal Component Analysis (PCA) further validated these differences, with tumor and normal samples forming distinct clusters along the principal components, reflecting the unique transcriptional landscapes of each group (**Fig. 1B, Supplementary Table 2**). A volcano plot highlighted the most notable genes, with the top 14 upregulated genes being Bystin like protein (BYSL), Solute Carrier Family 2 Member 13 (SLC2A13), Ectodermal Neural Cortex 1 (ENC1), Ajuba LIM Protein (AJUBA), Tumor Protein p53 Inducible Nuclear Protein 2 (TP53INP2), DEAD Box Helicase 56 (DDX56), Cbp/p300 Interacting Transactivator 2 (CITED2), PTEN Induced Kinase 1 (PINK1), Guanine Nucleotide Binding Protein G(I)/G(S)/G(O) Subunit Gamma 2 (GNG2), Semaphorin 6D (SEMA6D), Claudin 1 (CLDN1), Erythrocyte Membrane Protein Band 4.1 Like 3 (EPB41L3), Tetraspanin 7 (TSPAN7), ABI Family Member 3 Binding Protein (ABI3BP), and the top 27 downregulated genes including Lymphatic Vessel Endothelial Hyaluronan Receptor 1 (LYVE1), Phospholipid Phosphatase 1 (PLPP1), Nuclear Factor, Erythroid 2 Like 3 (NFE2L3), Electron Transfer Flavoprotein Dehydrogenase (ETFDH), Nuclear Receptor Subfamily 3 Group C Member 2 (NR3C2), Solute Carrier Organic Anion Transporter Family Member 4A1 (SLCO4A1), GTF2I Repeat Domain Containing 1 (GTF2IRD1), Twinkle mtDNA Helicase (TWNK), ETS Variant Transcription Factor 4 (ETV4), Transcription Factor 21(TCF21), Protein Phosphatase 2 Regulatory Subunit 3 Alpha (PPP2R3A), Sphingomyelin Phosphodiesterase 1 (SMPD1), Glycolipid Transfer Protein (GLTP), RuvB Like AAA ATPase 1 (RUVBL1), Purinergic Receptor P2Y1 (P2RY1), Thyroid Hormone Receptor Interactor 13 (TRIP13), Contactin 4 (CNTN4), Methylenetetrahydrofolate Dehydrogenase (NADP+ Dependent) 1 Like (MTHFD1L), Fibrinogen Like 2 (FGL2), Neuronal Growth Regulator 1 (NEGR1), Interleukin 6 Receptor (IL6R), Thiol Methyltransferase 1A (TMT1A), UDP-Glucose Pyrophosphorylase 2 (UGP2), Tribbles Pseudokinase 3 (TRIB3), Pleiotrophin (PTN), Guanine Nucleotide-Binding Protein G(I)/G(S)/G(O) Subunit Gamma-7 (GNG7), and Leukocyte Immunoglobulin-Like Receptor Subfamily B Member 5 (LILRB5). These genes exhibited highly significant (p-value < 0.05) (**Fig. 1C, Supplementary Table 2**). The differential expression patterns of these genes provide valuable insights into potential biomarkers and therapeutic targets in colon cancer.

**Figure 1.**
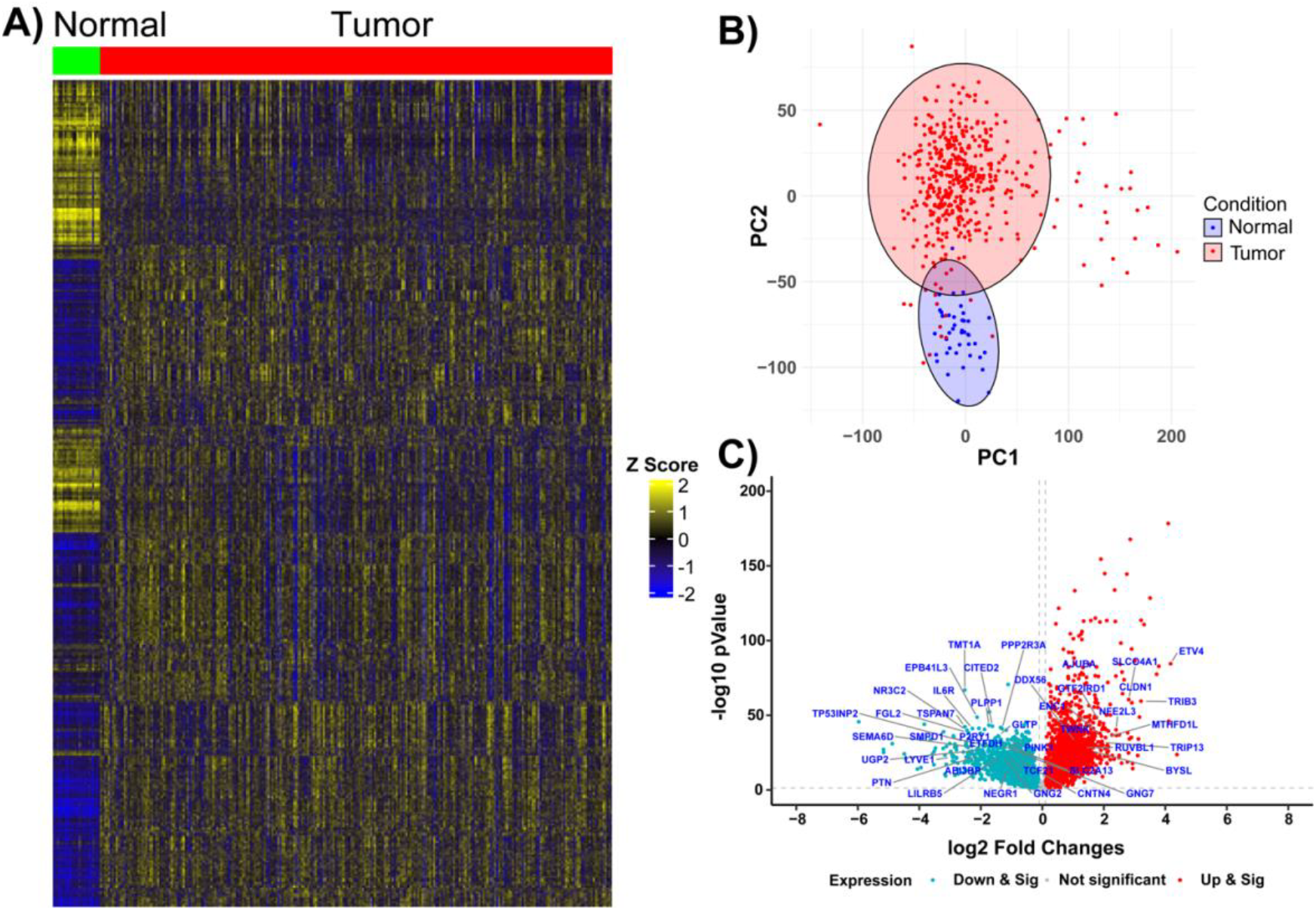
Differentially Expressed Genes in Tumor Tissue Samples of Colon Cancer A). The heatmap illustrates differentially expressed genes in tumor tissues (n = 444) compared to normal tissues (n = 41) from colon cancer patients. The gradient color scale represents increased expression (yellow) and decreased expression (blue). **B)**. The PCA plot depicts the clustering of tumor and normal tissue samples from colon cancer patients. **C)**. The volcano plot visualizes the distribution of differentially expressed genes between tumor and normal samples. The highlighted genes represent significantly altered genes identified using machine learning models-Random Forest (RF), Neural Network (NN), and Logistic Regression (LR)-all of which achieved AUC values exceeding 97%.

### Gene Set Enrichment Analysis (GSEA)

Gene Set Enrichment Analysis (GSEA) identified several hallmark pathways significantly enriched in tumor samples compared to healthy controls. The top 10 pathways included E2F targets, MYC targets V1, G2M checkpoint, MYC targets V2, MTORC1 signaling, adipogenesis, unfolded protein response, fatty acid metabolism, myogenesis, and DNA repair (**Fig. 2A, Supplementary Table 4**). The top-ranked pathway, E2F targets, involved critical cell cycle regulatory genes such as CDC25B, MYBL2, MYC, TRIP13, and UBE2S (**Fig. 2A**). These genes are important for cell cycle progression and are transcriptionally regulated by E2F transcription factors, highlighting their significant role in colon cancer pathogenesis. Enrichment plots of key pathways revealed a distinct concentration of altered genes at one end of the ranked gene list, supporting their biological significance in colon cancer progression (**Fig. 2B-2D**). The top three pathways-E2F targets, MYC targets V1, and G2M checkpoint-showed strong positive enrichment in tumor samples relative to normal controls, reinforcing their critical involvement in tumorigenesis (**Fig. 2B-2D**). These findings underscore the dysregulation of core processes such as cell cycle control, metabolic signaling, and stress responses in colon cancer. The enrichment of pathways like MTORC1 signaling and unfolded protein response points to metabolic rewiring and cellular stress adaptation as key features of tumor biology. Additionally, the dysregulation of fatty acid metabolism and adipogenesis suggests a potential link between lipid metabolism and tumor progression.

**Figure 2.**
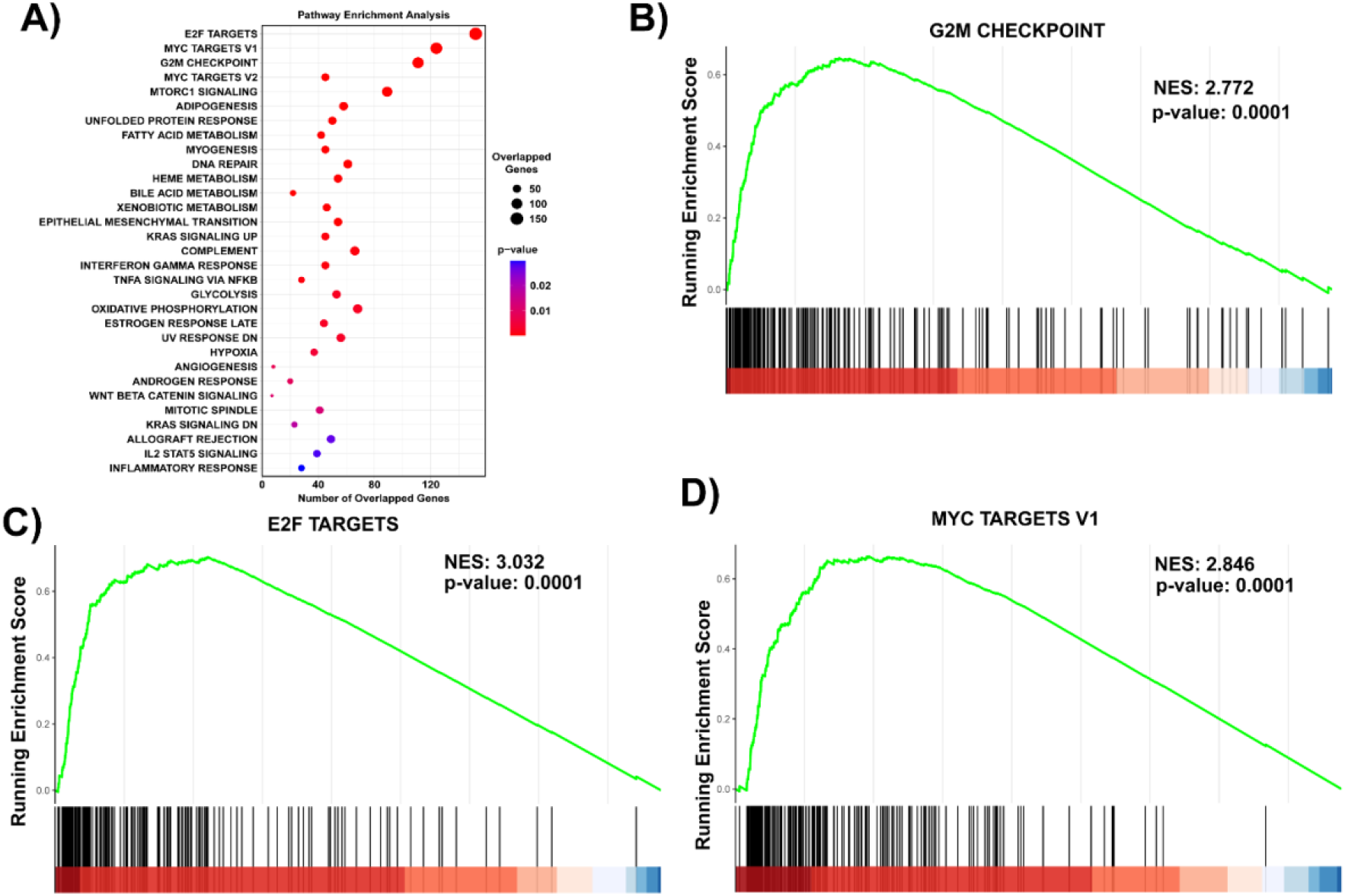
Gene Set Enrichment Analysis (GSEA) Pathway Analysis revealed the enriched biological pathways in colon cancer patients’ tumor samples. A). The dot plot represents the enriched pathways identified using GSEA. Pathways are ranked based on enrichment scores, with dot size corresponding to the number of overlapping genes and color indicating statistical significance (adjusted p-value). The enrichment plots display the top three significantly enriched pathways: G2M Checkpoints (B), E2F Targets (C), and MYC Targets (D). The green curve represents the running enrichment score, indicating the accumulation of gene hits as ranked by their differential expression. Vertical black lines denote the positions of pathway genes within the ranked gene list. The normalized enrichment score (NES) and p-values are displayed for each pathway.

### Machine Learning Analysis

The machine learning models, including Random Forest (RF), Neural Network (NN), and Logistic Regression (LR), achieved robust classification performance in distinguishing tumors from normal samples. The RF model demonstrated the highest accuracy (97%) and AUC (1.00), followed by NN (99%, AUC = 1.00) and LR (99%, AUC = 1.00). Area under curve (AUC > 0.97) analysis from the RF, NN, and LR model identified 41 top contributors’ genes to classification performance. These genes were consistently highlighted across models, indicating their potential as robust biomarkers and significantly altered in tumor samples compared to healthy samples in colon cancer (**Fig. 3, Supplementary Table 5 and 6**). The AUC-ROC curves demonstrate the models’ performance in distinguishing between classes, with the area under the curve (AUC) serving as a metric for classification accuracy. Higher AUC values indicate better model performance. The k-fold cross-validation approach ensures robustness and generalizability by partitioning the dataset into multiple subsets for training and validation (**Supplementary Fig. 1**). The top 3 genes (log2FC >3.00) of EPB41L3, TSPAN7, and ABI3BP genes were significantly increased, whereas another top 3 genes (log2FC < -2.5) of LYVE1, PLPP1, and NFE2L3 were significantly decreased in tumor samples compared to normal samples in colon cancer (**Fig. 3**).

**Figure 3.**
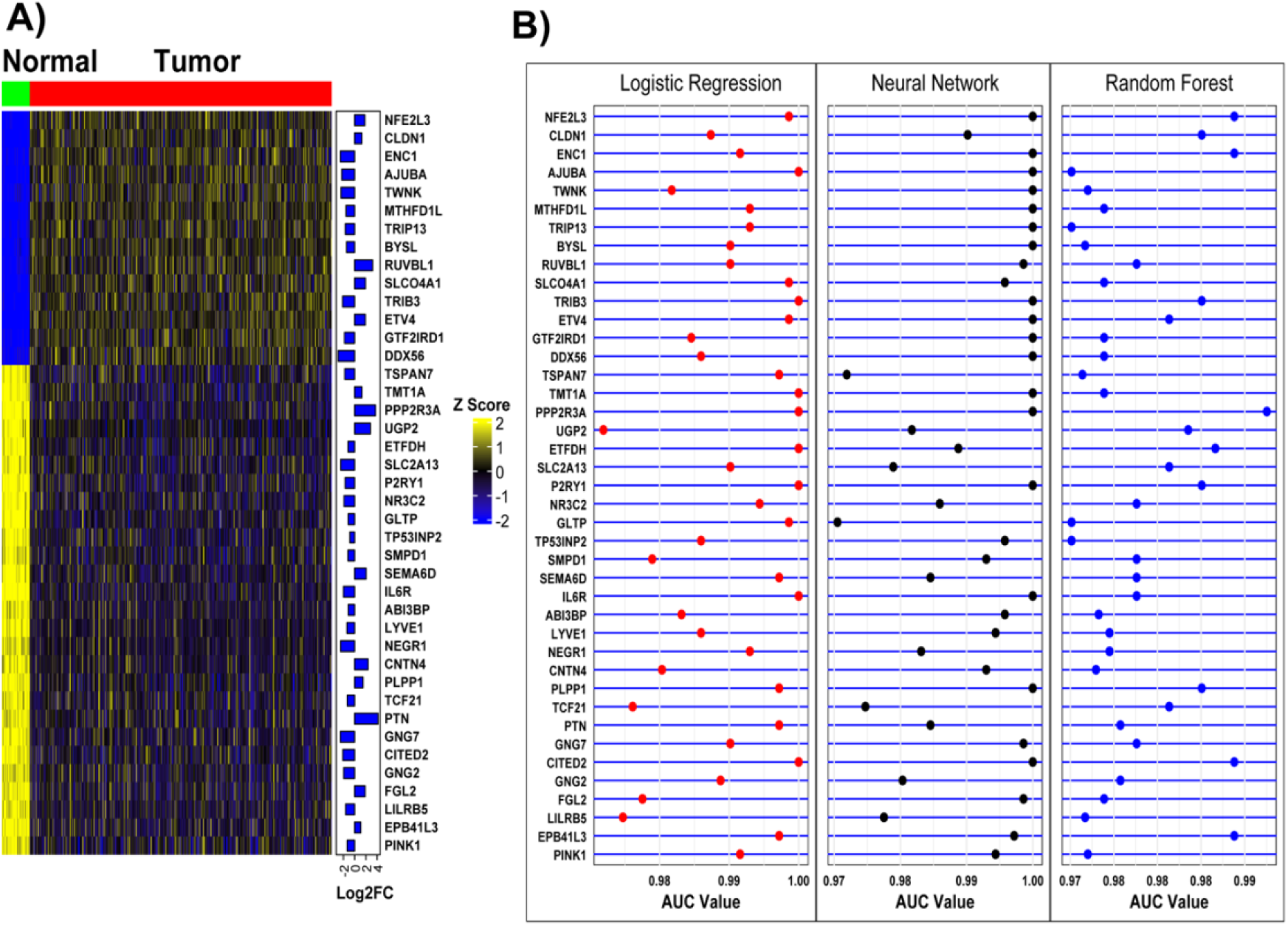
Machine Learning Models Classify Signature Genes in Tumor Patients. A). The heatmap displays differentially expressed genes and their fold changes in tumor tissues compared to normal tissues from colon cancer patients. These genes were identified using machine learning models-Random Forest (RF), Neural Network (NN), and Logistic Regression (LR)-all of which achieved AUC values exceeding 97%. **B)**. The dot plot illustrates the AUC distribution across the three machine learning models in tumor tissue samples, highlighting their classification performance.

### Identification of Potential Biomarkers

The integrative analysis, leveraging differential gene expression, Gene Set Enrichment Analysis (GSEA), and machine learning approaches, identified a subset of genes with high predictive potential in colon cancer. The top three upregulated genes were EPB41L3, TSPAN7, and ABI3BP, which exhibited significantly increased expression in tumor samples compared to normal samples (**Fig. 3**). These genes are likely involved in tumor progression and could serve as potential targets for therapeutic intervention. Conversely, the top three downregulated genes were LYVE1, PLPP1, and NFE2L3, which were significantly suppressed in tumor samples (**Fig. 3**). Their decreased expression may indicate disruption in pathways critical for maintaining normal cellular homeostasis and immune response. These findings underscore the potential of these genes as robust biomarkers for distinguishing tumors from normal samples. Furthermore, their consistent identification across multiple analytical approaches lays a strong foundation for subsequent validation studies. Ultimately, these biomarkers hold promise for advancing colon cancer diagnostics and therapeutics, paving the way for personalized medicine strategies.

## Discussion

This study demonstrates the power of integrating transcriptomics and machine learning to uncover robust biomarkers and pathways in colon cancer. Our findings reveal distinct transcriptional and pathway-level alterations that differentiate tumors from normal samples, providing critical insights into colon cancer biology. By combining differential gene expression analysis, GSEA, and machine learning approaches, we uncovered distinct transcriptional and pathway-level alterations that differentiate tumors from normal samples. These findings not only deepen our understanding of colon cancer biology but also lay a foundation for the development of diagnostic and therapeutic strategies.

The differential gene expression analysis revealed a substantial number of genes with altered expression in tumor samples, including 657 upregulated and 8,566 downregulated genes. Among these, EPB41L3, TSPAN7, and ABI3BP emerged as the most upregulated genes (log2FC > 3.00), suggesting their potential involvement in tumor progression. These genes have been implicated in cellular adhesion, signaling, and modulation of the tumor microenvironment, all critical processes in cancer^10^. Conversely, the top three downregulated genes, LYVE1, PLPP1, and NFE2L3 (log2FC < -2.5), suggest disrupted immune signaling and lipid metabolism in tumor tissues. For instance, LYVE1 is linked to lymphatic vessel integrity and immune regulation^9^, and its suppression may contribute to immune evasion. Similarly, PLPP1 and NFE2L3, associated with lipid metabolism and oxidative stress^8^, underscore the metabolic vulnerabilities of tumor cells.

Gene Set Enrichment Analysis (GSEA) provided additional insights into the pathways disrupted in colon cancer. Key pathways, including E2F targets, MYC targets, and G2M checkpoints, were significantly enriched in tumor samples. These findings emphasize dysregulated cell cycle control, proliferation, and metabolic rewiring as hallmarks of colon cancer^7^. The identification of pathways such as MTORC1 signaling and unfolded protein response highlights the ability of colon cancer cells to adapt to metabolic stress and optimize survival in nutrient-limited environments^11^. Enrichment of pathways like fatty acid metabolism and adipogenesis further underscores the interplay between lipid metabolism and tumor progression.

Machine learning analyses reinforced the robustness of the identified biomarkers. The Random Forest, Neural Network, and Logistic Regression models achieved high classification accuracy (97-99%) and AUC values (1.00), demonstrating their predictive power in distinguishing tumors from normal samples. Importantly, identifying key genes across models consistently underscores their translational relevance. Genes such as EPB41L3^19^, TSPAN7^20^, and ABI3BP^21-24^, as well as LYVE1^25-27^, PLPP1^28^, and NFE2L3^29-32^, were consistently highlighted as significant contributors to classification performance. These results confirm the utility of integrating transcriptomics with machine learning for biomarker discovery^11^.

The observed transcriptional and pathway alterations align with previous studies while also providing novel insights into the molecular underpinnings of colon cancer^33^. For instance, the enrichment of E2F and MYC target pathways supports the role of dysregulated transcriptional networks in driving tumor growth^9, 34, 35^. Additionally, the suppression of immune-related genes, such as LYVE1^25-27^, highlights potential mechanisms of immune evasion, which may be critical for tumor survival and progression^36^.

While this study provides valuable insights, there are limitations that warrant further investigation. Validation in independent cohorts and experimental models is essential to confirm the identified biomarkers and pathways. Additionally, functional studies are needed to elucidate the precise roles of these genes and pathways in colon cancer pathogenesis. Integrating additional omics datasets, such as proteomics or metabolomics, could offer a more comprehensive understanding of tumor biology and uncover additional therapeutic opportunities^10^.

In conclusion, our integrative approach identifies signature genes and pathways in colon cancer, offering promising candidates for diagnostic and therapeutic development. Future studies should focus on validating these biomarkers in independent cohorts and exploring their functional roles to advance precision medicine in colon cancer. The identified genes and pathways provide promising candidates for diagnostic and therapeutic applications, paving the way for precision medicine strategies. Future studies focusing on experimental validation and clinical translation will be crucial for leveraging these findings to improve patient outcomes in colon cancer.

## Author Contributions

Conceptualization: M. A. H, and M. S.I; Methodology: M. A. H; Data Collection: M. A. H; Manuscript Preparation: M. A. H, and M. S.I; Writing-review and editing: M. A. H, and M. S.I; All authors have read and agreed to the published version of the manuscript.

## Ethical approval

Not applicable

---

**Supplementary Figure 1.**
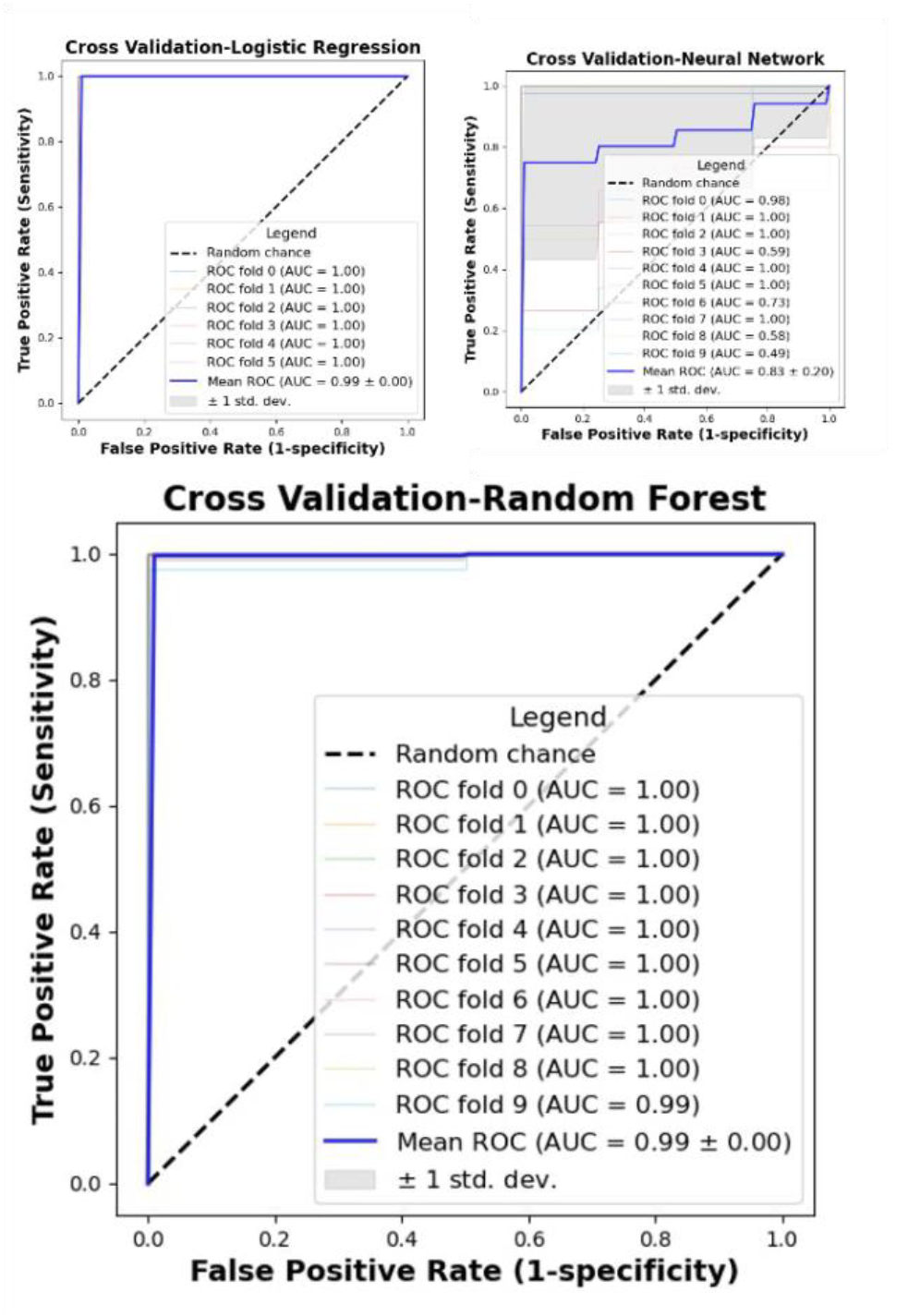
AUC-ROC Curves of k-Fold Cross-Validation for Machine Learning Models. The figure illustrates the Area Under the Receiver Operating Characteristic (AUC-ROC) curves for three machine learning models-Logistic Regression (LR), Neural Network (NN), and Random Forest (RF)-evaluated using k-fold cross-validation. The models were trained and tested on the dataset with the following configurations: Logistic Regression (LR): Utilized 5 splits for k-fold cross-validation. Neural Network (NN) and Random Forest (RF): Employed 9 splits for k-fold cross-validation. Random Forest (RF): Also used 9 splits for k-fold cross-validation.

