## Supplementary Table 6 for "Integrative Machine Learning Reveals Potential Signature Genes Using Transcriptomics in Colon Cancer"

### Slide 1
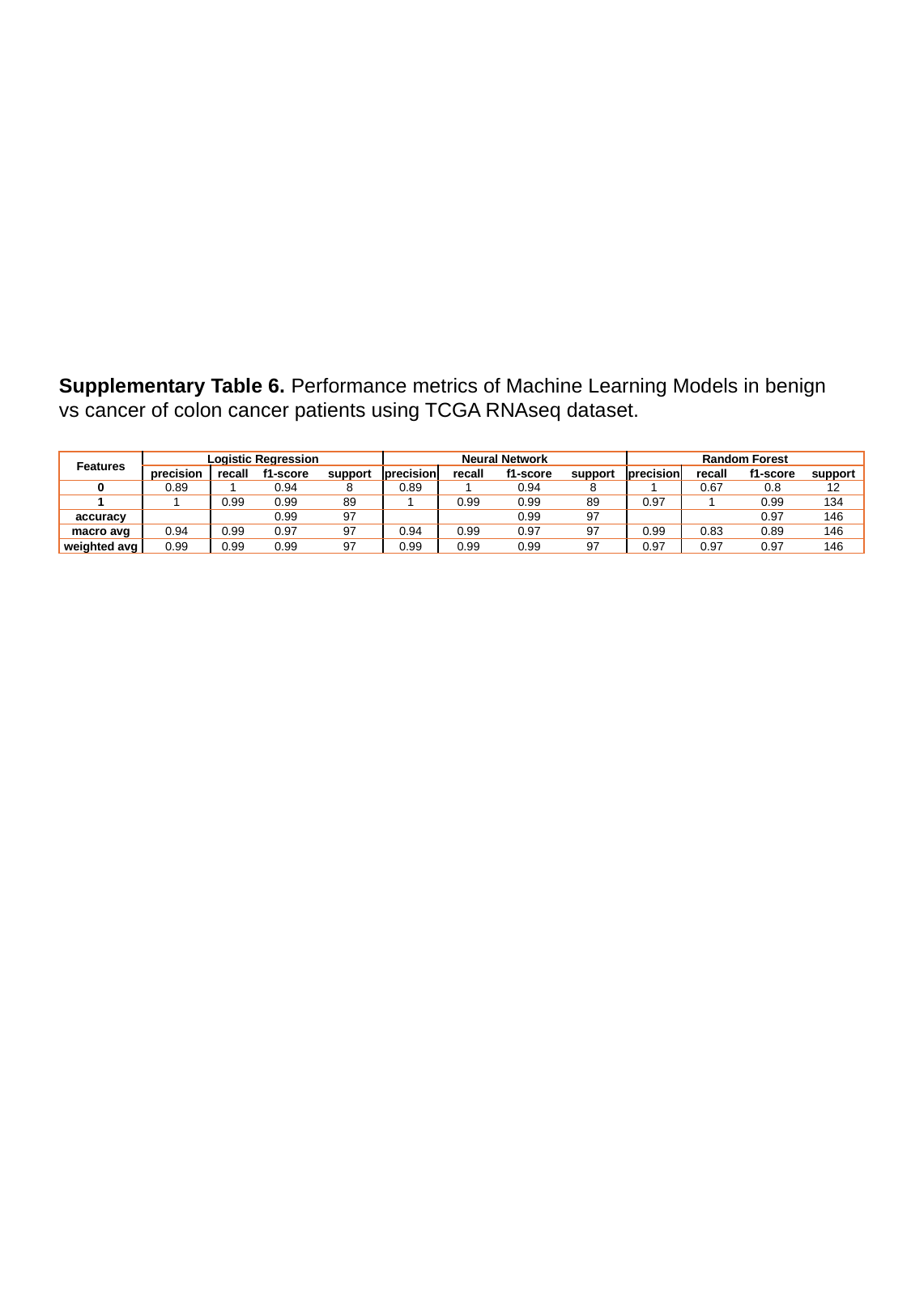

Supplementary Table 6. Performance metrics of Machine Learning Models in benign vs cancer of colon cancer patients using TCGA RNAseq dataset.
| Features | Logistic Regression | | | | Neural Network | | | | Random Forest | | | |
| --- | --- | --- | --- | --- | --- | --- | --- | --- | --- | --- | --- | --- |
| | precision | recall | f1-score | support | precision | recall | f1-score | support | precision | recall | f1-score | support |
| 0 | 0.89 | 1 | 0.94 | 8 | 0.89 | 1 | 0.94 | 8 | 1 | 0.67 | 0.8 | 12 |
| 1 | 1 | 0.99 | 0.99 | 89 | 1 | 0.99 | 0.99 | 89 | 0.97 | 1 | 0.99 | 134 |
| accuracy | | | 0.99 | 97 | | | 0.99 | 97 | | | 0.97 | 146 |
| macro avg | 0.94 | 0.99 | 0.97 | 97 | 0.94 | 0.99 | 0.97 | 97 | 0.99 | 0.83 | 0.89 | 146 |
| weighted avg | 0.99 | 0.99 | 0.99 | 97 | 0.99 | 0.99 | 0.99 | 97 | 0.97 | 0.97 | 0.97 | 146 |
